# Accelerated physiology and increased energy expenditure in animals and humans with mitochondrial defects: A meta-analysis

**DOI:** 10.1101/2023.09.09.556754

**Authors:** Alexander J. Sercel, Gabriel Sturm, Evan D. Shaulson, Dympna Gallagher, Marie-Pierre St-Onge, Christopher P. Kempes, Herman Pontzer, Michio Hirano, Martin Picard

## Abstract

Mitochondria are key energy transforming organelles in mammalian cells. However, how defects in oxidative phosphorylation (OxPhos) and other mitochondrial functions influence whole-body energy expenditure (EE) has not been rigorously studied. Cellular and organismal responses to OxPhos defects likely involve a combination of functional *downregulation* to conserve energy and compensatory *upregulation* of stress responses. If the energy cost of compensatory responses exceeds the potential energy savings of functional downregulation, as recent work suggests, the result would be an increase in total EE. To address the hypothesis that OxPhos defects increase the energetic cost of living, we performed a meta-analysis of available studies reporting EE in animal models with mitochondrial gene defects. Of all reported experimental conditions (n = 91, from 29 studies), 51% reported a >10% elevation in EE relative to control animals, compared to 11% reporting <10% reduction in EE (p<0.0001, Chi-square). Of the experimental conditions where locomotor activity was also quantified, 39% showed that OxPhos-deficient animals had elevated EE despite reduced locomotor activity, which would be expected to decrease EE. To extend this finding in humans, we re-analyzed a high-quality clinical and multi-omics dataset (Sharma et al. 2021) of mitochondrial disease patients with the m.3243A>G mutation. This analysis similarly indicates an upregulation of energetically costly physiological, immune, and metabolic parameters in people with OxPhos deficiency. These results suggest that animals and humans with mitochondrial defects must expend more energy to sustain life, a state clinically called *hypermetabolism*. High-quality human energetics studies are needed to understand the magnitude, mechanisms, and modifiability of hypermetabolism in mitochondrial disorders.

## Introduction

In mammals, the Krebs cycle and respiratory chain within mitochondria transform chemical energy to generate an electrochemical potential that powers dozens of key cellular processes, including ATP synthesis by oxidative phosphorylation (OxPhos)^1^. Failure of OxPhos and loss of mitochondrial membrane potential leads to death within seconds to minutes, illustrating the vital role of this process. Moreover, mitochondria also regulate information flow^2^. Mitochondria contain dozens of sensing mechanisms that tune their anabolic and catabolic activities and produce molecular signals that reach the nucleus to alter cellular activities and the systemic circulation to regulate whole-body physiology^2^. Although mitochondria are key to processing energy and information, the energetic cost of impaired mitochondrial OxPhos to an organism remains unclear.

Hundreds of characterized nuclear or mitochondrial DNA (mtDNA) mutations causing OxPhos defects result in a diverse range of phenotypes called mitochondrial diseases^3^. Individuals with a mitochondrial disease typically present with short stature and low BMI, experience debilitating fatigue and exercise intolerance, and display heterogenous multi-organ symptoms, which in their most severe form, manifest as multi-organ failure and premature death^4-6^. While the exact mechanisms of pathogenesis leading from mutation to clinical presentation of mitochondrial disease remains unclear, much is known about the impact of OxPhos dysfunction at the cell, tissue, and systems level in patients.

*Cells* with OxPhos defects shift to glycolytic metabolism^7^, increase mitochondrial biogenesis, accumulate mitochondrial mass and excess mtDNA copies^8,9^, activate the integrated stress response (ISR)^7,10-12^, secrete signaling factors including the classic metabokines fibroblast growth factor 21 (FGF21) and growth differentiation factor 15 (GDF15)^7^, and can exhibit DNA instability^7^. At the *+ssue* level, mitochondrial disease patients and mouse models of OxPhos defects show elevated levels of circulating metabokines^13^, catecholamines^7^, and inflammatory cytokines^14^, exhibit hypercapillarization (i.e., excessive angiogenesis) selectively around OxPhos-defective cells^15,16^, and may have impaired immunity^17,18^. Clinically, patients suffer from *system* level effects, such as lactic acidosis^19^, increased resting heart rate^7^, hyperkinetic cardiorespiratory responses to mild physical activity^20,21^, and report greater stress and anxiety^22^. Thus far, there is no unifying framework accounting for the complex clinical presentation of mitochondrial diseases. How do loss-of-function OxPhos defects produce this constellation of apparent gain-of-function symptoms?

Despite the key roles that mitochondria play in energy sensing and processing, we do not know how defects in OxPhos and other mitochondrial functions impact the energetic requirements of organisms. Furthermore, the role of energy homeostasis in the symptomatic progression of mitochondrial disease is not well understood. Studies measuring the basal energetic requirements of patients with mitochondrial disease are limited^23,24^. The community’s research efforts have primarily focused on elucidating the molecular drivers of mitochondrial diseases, without an eye towards the potential influence of energy consumption rates. To address this gap, here we present a meta-analysis of energy expenditure (EE) in animal models with genetic mutations in mitochondrial genes. We also re-analyze the most comprehensive clinical and multi-omics dataset in a cohort of human patients with the most common pathogenic mtDNA mutation^25^. Together, these data suggest that mitochondrial OxPhos defects consistently increase whole-body energetic requirements, which may have important implications for the etiology and potential treatments of mitochondrial diseases.

## Results

### Energy expenditure measurements in animal models of mitochondrial mutations

To investigate the effect of mitochondrial defects on whole-body EE, we searched the Medline database through PubMed for studies quantifying EE in animal models with genetic mutation, knock-out, or overexpression of mitochondrial genes. Genes targeted include OxPhos subunits and assembly factors, as well as non-OxPhos genes involved in mitochondrial dynamics, transport, mtDNA replication, or quality control. The resulting studies (n=29) are listed in Table 1 and organized by genetic manipulation. Available data include species, targeted tissue (in the case of tissue-specific models), age, activated cellular stress signaling (where available), method to quantify and normalize whole-body EE, and the specific experimental conditions (e.g., time of day) used for comparison to control animals. We extracted data from a total of 91 experimental conditions and report the calculated difference in EE relative to the control group in each study (EE % of control), along with ambulatory activity or physical activity levels also relative to the control group of each study.

**Table 1.**
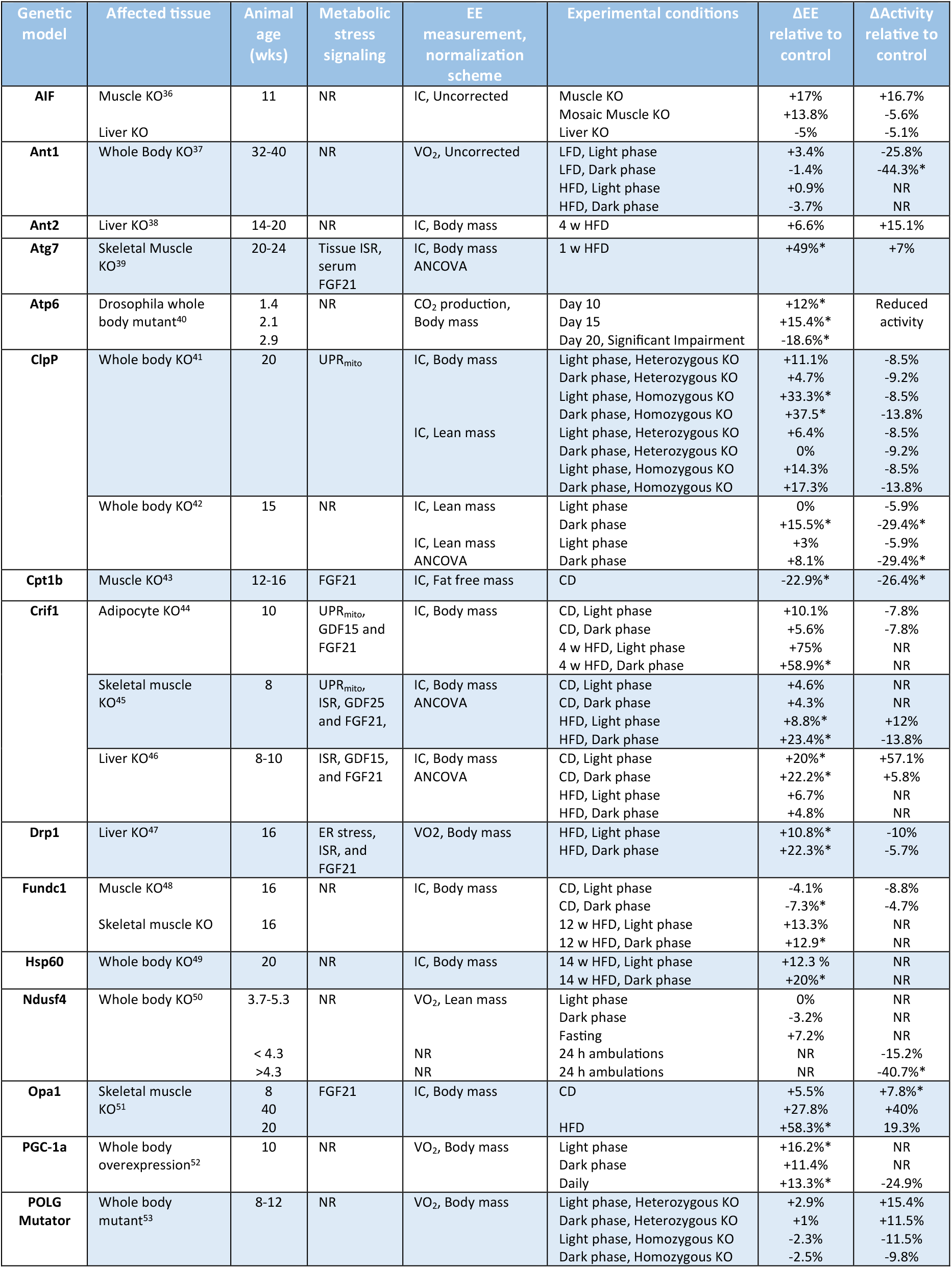
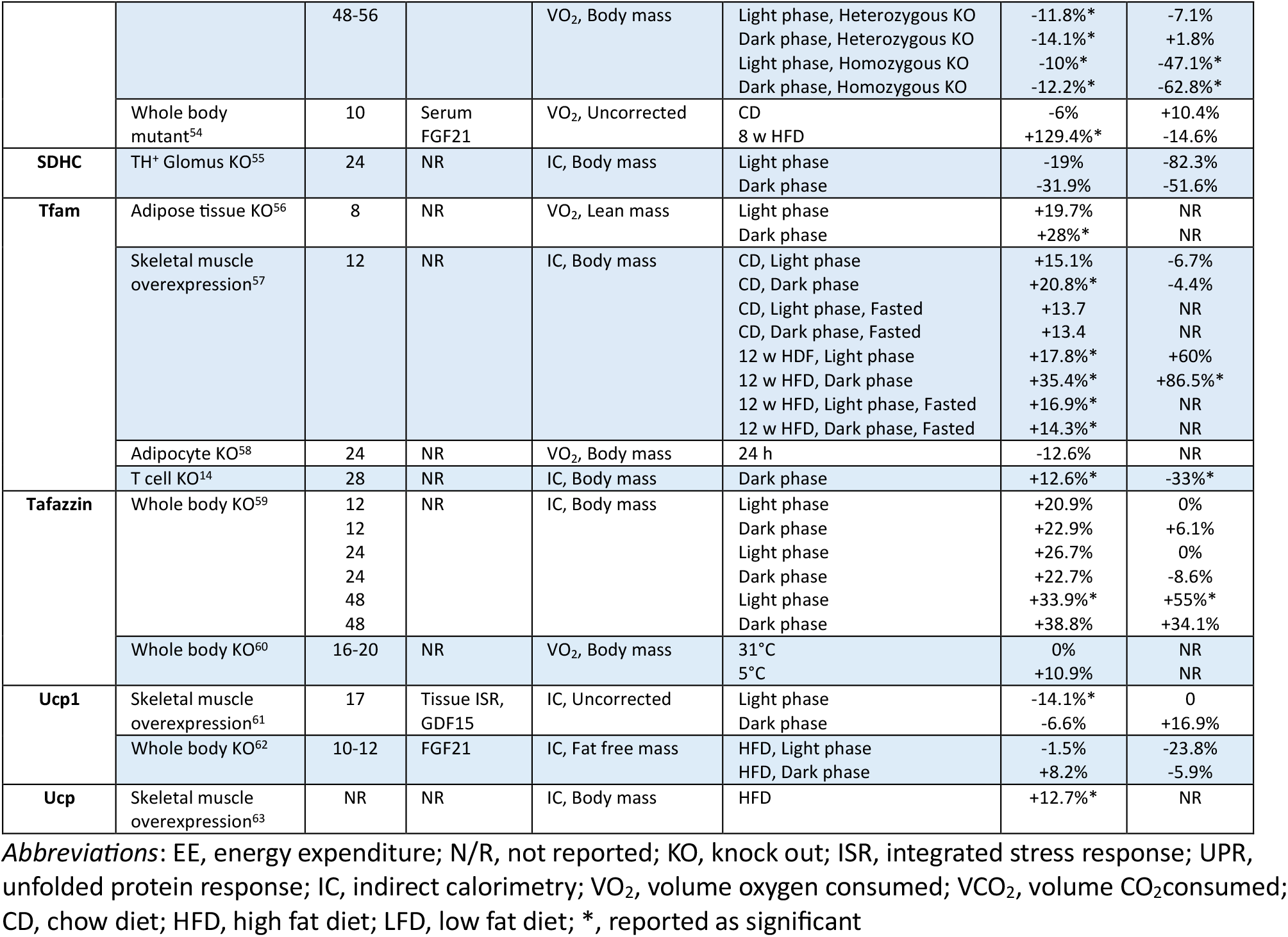
Evidence of hypermetabolism in animal models of mitochondrial OxPhos defects. Animal studies, few of which have used the appropriate normalization approaches and none of which have specifically examined hypermetabolism, are summarized with their effect sizes. A re-analysis of available data, taking into consideration changes in locomotor activity, points to elevated energy expenditure (EE) independent of elevated physical activity as a common feature in OxPhos-deficient animals.

To interpret the meta-analysis results, we projected the EE and locomotor activity data into a 2-dimensional space (Figure 1a). All things being equal, animals who move more should expend more energy, whereas animals who are less active should have lower EE^26,27^. Therefore, accounting for differences in physical activity levels is critical to establish if OxPhos-deficient animals exhibit differences in resting EE. Resting EE on the other hand, reflects the energetic cost of somatic maintenance and other intrinsic (physiological, cellular, molecular) processes. In the 2-dimensional space, we define a conservative “hypermetabolic” quadrant (*bottom right*) to describe animals with elevated EE and reduced locomotor activity. The inverse phenomenon, where animals show reduced EE despite elevated locomotor activity, is termed “hypometabolic” (*top left*). We refer to experimental conditions with roughly proportional increases or reductions in EE and locomotor activity as “normometabolic”, noting that false-negatives (animals with severely reduced activity but only mildly reduced EE; or animals with greatly elevated EE but modest elevation in activity, classified as normometabolic) may occur in this simple system.

**Figure 1.**
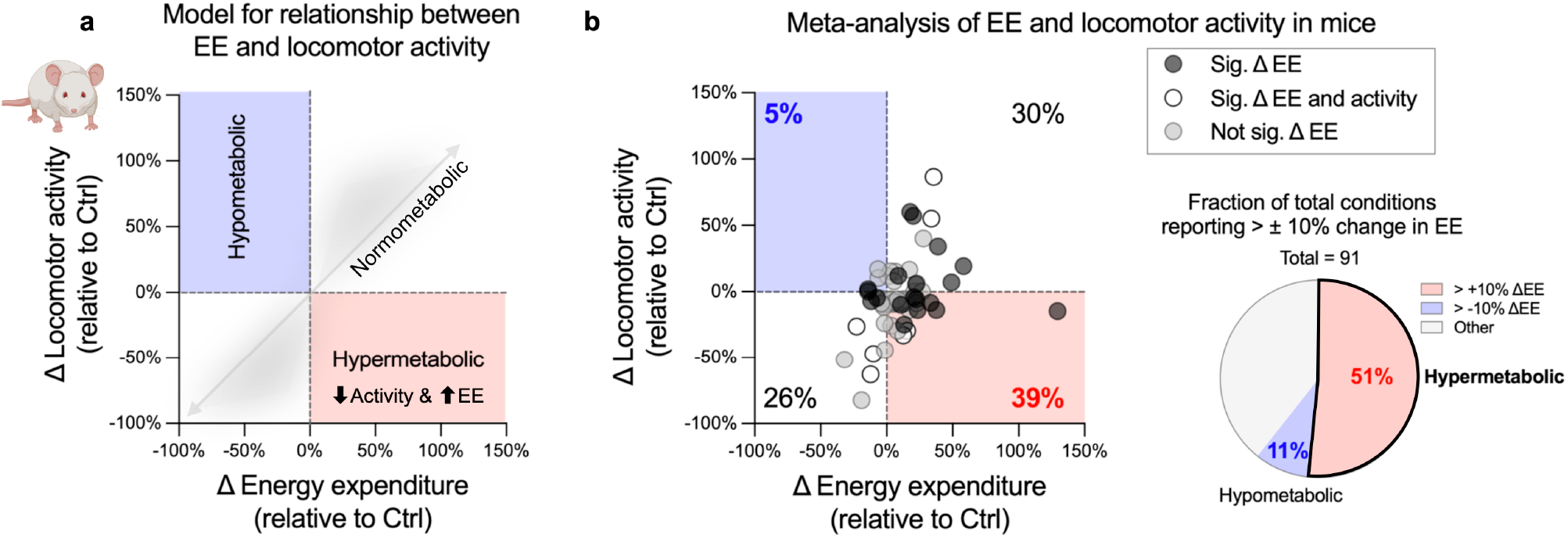
Meta-analysis of OxPhos-deficient animal models reveals a signature of elevated energy expenditure not hypometabolism and ATP deficit. **a**) Visualization of the three metabolic modes – hypometabolism, normometabolism, and hypermetabolism – that animal models of OxPhos dysfunction can theoretically adopt as plotted on the axes of energy expenditure and locomotor activity. **b**) Meta-reanalysis of published reports (n = 19) comparing changes in EE (as normalized in the original manuscripts) and locomotor activity of OxPhos-deficient animals (source data in Table 1). The percentages of total observations (n = 52) in each of the four quadrants is noted and significantly different from the null hypothesis tested by chi-square, p<0.0009. The pie chart shows the proportion of all measurements of OxPhos-deficient animals (n = 78) with EE greater than 10% or less than -10% of control values. The distribution observed is significantly different from the null hypotheses tested by chi-square, p<0.002.

Across all experimental conditions with both EE and locomotor activity (n=61), 39% showed evidence of hypermetabolism in OxPhos-deficient animals, significantly more than expected by chance (p<0.0015, Chi-square). In contrast, 5% showed hypometabolism, and 56% showed normometabolism. The number of studies in the “normometabolic” category is likely an overestimate owing to the (often unmeasured) reduced vigor and physical activity of OxPhos-deficient animals with similar EE as control animals, despite significantly reduced activity-related EE.

To examine these data using a different approach, we quantified the fraction of all experimental conditions reporting differences in EE relative to control (n=91), where the group difference is greater than ± 10% (as a conservative estimate), irrespective of activity levels. There were 11% of experimental conditions reducing EE by 10% or more, whereas 4.6 times more experimental conditions, or 51% of studies, reported a >10% higher EE (p<0.0001, Chi-square). 38% of experimental conditions were within ±10% of control, likely within measurement variability. Thus, the majority of animal studies indicate that OxPhos-deficient animals expend more energy at rest than their OxPhos-normal counterparts.

### Bias towards upregulation of metabolites and proteins in humans with OxPhos defects

To extend the meta-analyses of EE in animals models (above, Figure 1) and clinical studies^7^, we re-analyzed the most comprehensive clinical and multi-omics biomarker study in individuals with the m.3243A>G mtDNA mutation, reported by Sharma et al., 2021^25^. This mutation is the most common pathogenic mtDNA defect in the human population^28^. It causes a variable degree of disease severity depending, among other factors, on the mutation load or heteroplasmy (% mutant). It in its most severe form, the m.3243A>G mtDNA mutation causes mitochondrial encephalopathy, lactic acidosis, and stroke-like episodes (MELAS), which is associated with a 3-4-decade reduction in lifespan^29^.

We extend the conceptual model of hypo-, normo-, and hypermetabolism to a framework in which rates of molecular processes or levels of biomarkers in mitochondrial disease patients are reduced, equal to, or greater than in controls. We term these states hypoactivity, normoactivity, and hyperactivity, respectively. We reason that if the symptoms of mitochondrial disease were driven by a general slowing or reduction of molecular processes among cells and organs (as one may intuitively assume from the energy shortage), we would expect to see evidence of *hypoactivity* in OxPhos-deficient individuals. Conversely, if as in the cellular and animal systems, OxPhos-deficient cells and physiological systems expend excess energy to mount compensatory responses, we expect to observe evidence of physiological *hyperactivity* in people with mitochondrial diseases.

With this model in mind, we quantified the balance of differentially represented plasma proteins and metabolites measured by discovery-based platforms (total n=1,310 proteins, 376 known metabolites) between healthy controls and individuals with the m.3243A>G mutation. Here the null hypothesis is that an equal number of analytes would randomly be either down-or upregulated in mitochondrial diseases. In contrast, data from Sharma et al. identified 10% more upregulated proteins (686 in patients vs 624 in controls, n.s., Chi square) in patients relative to controls, and 55% more upregulated metabolites (228 vs 147, p<0.0001, Chi-square) (Figure 2a). Focusing only on the 23 statistically significant metabolites that were differentially represented between the two groups, 91.3% were upregulated (p<0.0001, compared to the 50:50 null hypothesis) in patients with mitochondrial disease. Consistent with the hypersecretion previously reported in cells with genetic or pharmacologically-induced OxPhos defects^7^, the excess abundance of both circulating proteins and metabolites paints a picture of physiological *hyperactivity*.

**Figure 2.**
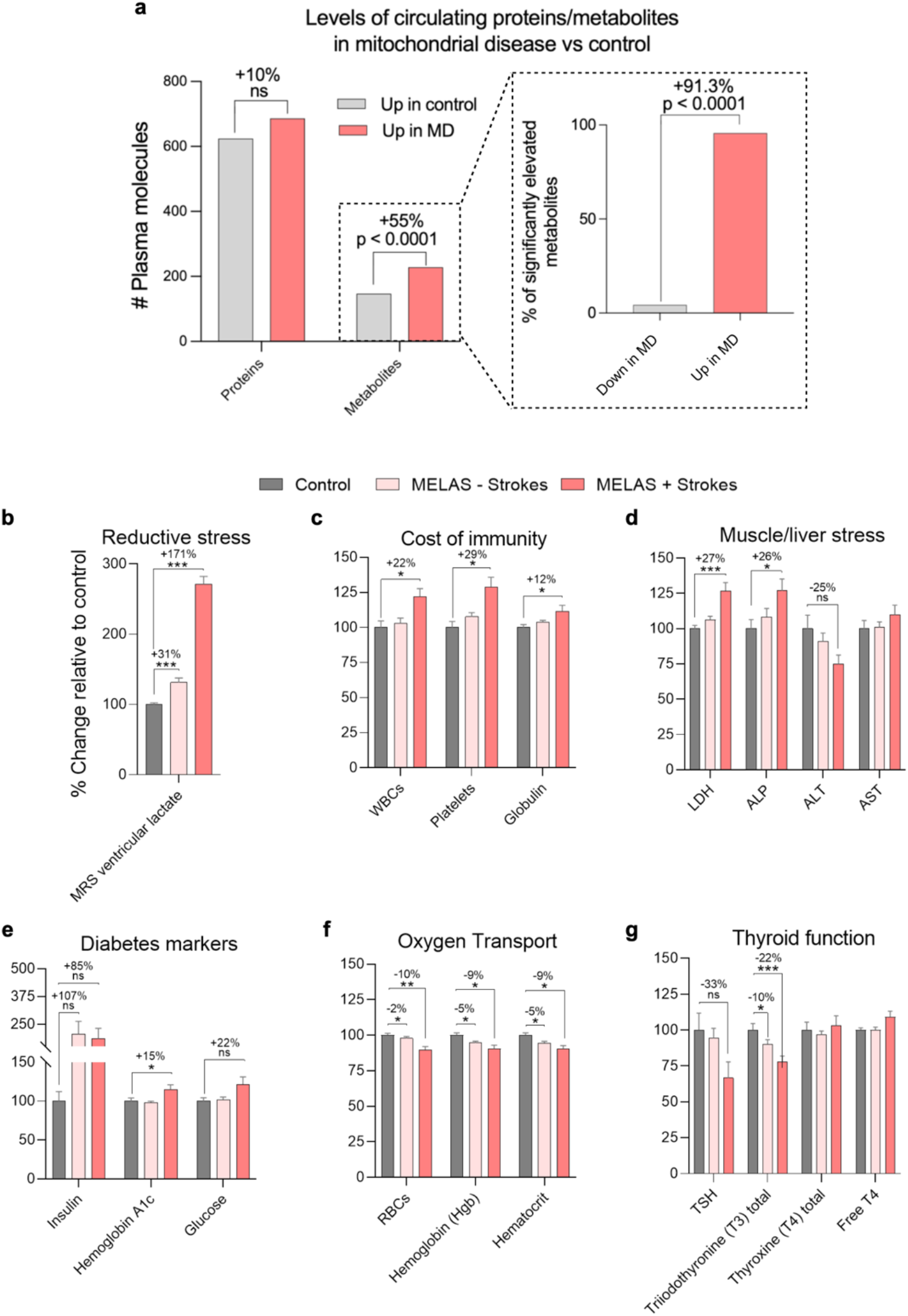
Re-analysis of multi-omics and clinical results from patients with mitochondrial disease reveal multi-system recalibration and physiological hyperactivity. **a**) Excess of upregulated plasma proteins and metabolites in patients with mitochondrial encephalopathy, lactic acidosis, and stroke like episodes (MELAS) vs controls relative to downregulated molecules. Data from Sharma et al. (2021), p values from chi square, n = 16-20 MELAS, 24-32 controls. **b-g**) Patients with OxPhos defects show the expected elevation in lactate reflecting reductive stress and evidence of physiological hyperactivity. They have an elevated number of circulating bone marrow-derived immune cells, elevated markers of muscle and liver damage, and elevated glucose and glucose-regulating insulin. In contrast and reflecting the reduced ability to consume oxygen, OxPhos-deficient patients downregulate oxygen transport and hypermetabolism-promoting thyroid function, possibly as an attempt to decrease whole-body EE. MELAS patients with strokes (severe, n = 13), MELAS patients without strokes (less severe, n = 76), and controls (n = 32). * P < 0.05, ** P < 0.01, *** P < 0.001, Wilcoxon rank-sum test. Abbreviations: *EE*, energy expenditure; *MD*, mitochondrial disease; *MRS*, magnetic resonance spectroscopy; *WBC*, white blood cells; *LDH*, lactate dehydrogenase; *ALP*, alkaline phosphatase; *ALT*, alanine transaminase; *AST*, aspartate transferase; *Hemoglobin A1c*; glycated hemoglobin; *RBCs*, red blood cells; *TSH*, thyroid stimulating hormone.

### Clinical analytics of MELAS suggests physiological hyperactivity

The study by Sharma et al. also quantified several standard laboratory analytes and classic blood biomarkers that reflect the activity of major physiological systems. This data is presented for three groups: i) healthy controls (n=32), ii) individuals who carry the m.3243A>G mutation but who do not have stroke-like episodes (“mild” mitochondrial disease, n=76), and iii) individuals with the same mutation but with MELAS (“severe” mitochondrial disease, n=13). As expected, there was a significant symptom-related increase in brain ventricular lactate in patients (+31% without stroke-like episodes, +171% with stroke-like episodes) (Figure 2b), congruent with the presence of OxPhos deficiency and systemic lactic acidosis.

To test our central hypothesis and gain insight into what might be contributing to hypermetabolism in patients with OxPhos defects^7^, we focused our analysis of the clinical biomarkers that either reflect energy-consuming physiological processes, or that are primarily energy-regulating in nature. The immune system incurs significant energetic costs required to build, maintain, and activate the immune repertoire^30,31^. Interestingly, OxPhos-deficient patients with MELAS had significantly *higher* total white blood cells (+22%), platelet count (+29%), and plasma globulin level (+12%) (Figure 2c). Because immune cell maintenance and activity are energetically costly, this relatively mild leukocytosis is expected to increase whole-body energy demands. Moreover, MELAS patients also showed significantly higher levels of the muscle and liver stress markers lactate dehydrogenase (+27%) and alkaline phosphatase (+26%) (Figure 2d). These enzymes are not only energetically costly to synthesize, but may also mark the presence of cellular damage requiring energy-dependent repair processes in the source tissue(s).

In relation to energy-regulating and delivering systems, patients showed a non-significant but elevated level of plasma insulin (+107% without stroke-like episodes, +85% with stroke-like episodes), and severe MELAS patients showed significantly elevated hemoglobin A1c (+15%) and glucose (+22%) levels, indicating physiological metabolic stress (Figure 2e). Physiological hyperglycemia may be required to support hypermetabolism, but may incur additional energetic costs to degrade, resynthesize, and compensate for the accumulated glycated extracellular and intracellular proteins.

Consistent with impaired oxygen extraction and demand in OxPhos-deficient tissues, MELAS patients had lower levels of oxygen transport markers including total red blood cell content (RBCs, -10%), hemoglobin (-9%), and hematocrit (-9%) (Figure 2f).

Lastly, consistent with a state of hypermetabolism of non-thyroidal origin^32,33^, circulating levels of thyroid hormones were reduced in patients, including 33% lower thyroid stimulating hormone (TSH) and 22% lower total triiodothyronine (T_3_) in MELAS patients (Figure 2g). The downregulation of thyroid hormone synthesis originating at the level of TSH largely rules out primary thyroid gland dysfunction, instead suggesting a physiological strategy aiming to decrease the output of the naturally hypermetabolism-stimulating T_3_ and T_4_ hormones.

## Discussion

Together, the meta-analysis of animal studies and re-analysis of clinical results reveal a physiological signature consistent with elevated EE and physiological hyperactivity in animals and humans with molecular mitochondrial defects. Our meta-analysis shows increased whole-body EE in the absence of increased physical activity, suggesting that mutations in mitochondrial genes elevate the energetic cost of life in tissues at rest. Additionally, the re-analysis of clinical blood measurements from Sharma et al. highlights multiple markers of hyperactive physiology, and of the potential whole-body response contributing to hypermetabolism in individuals with OxPhos defects. In particular, the elevated WBC count, antibody levels, cytokines/metabokines, chronic hyperglycemia, and the downregulation of metabolism-stimulating thyroid hormone signaling, paint a rather coherent physiological picture of stress-evoked hypermetabolism.

These results align with reports of mitochondrial disease patients showing high levels of fatigue and avoiding physical activity due to exercise intolerance, accompanied by elevations in both resting heart rate and resting oxygen consumption^7^. These findings also are mirrored by *in vitro* studies of OxPhos-deficient patient-derived cells and of normal cells with acutely-induced OxPhos deficiency, both of which mount similar integrated stress responses, exhibit hypersecretory phenotypes, and age at an accelerated pace^7,10^. Thus, our analysis suggests that cells and animals do not respond to OxPhos defects by slowing down or quieting their physiology, but rather actively mobilize energy-consuming integrated responses that likely contribute to hypermetabolism and accelerate disease presentation and progression (Figure 3).

**Figure 3.**
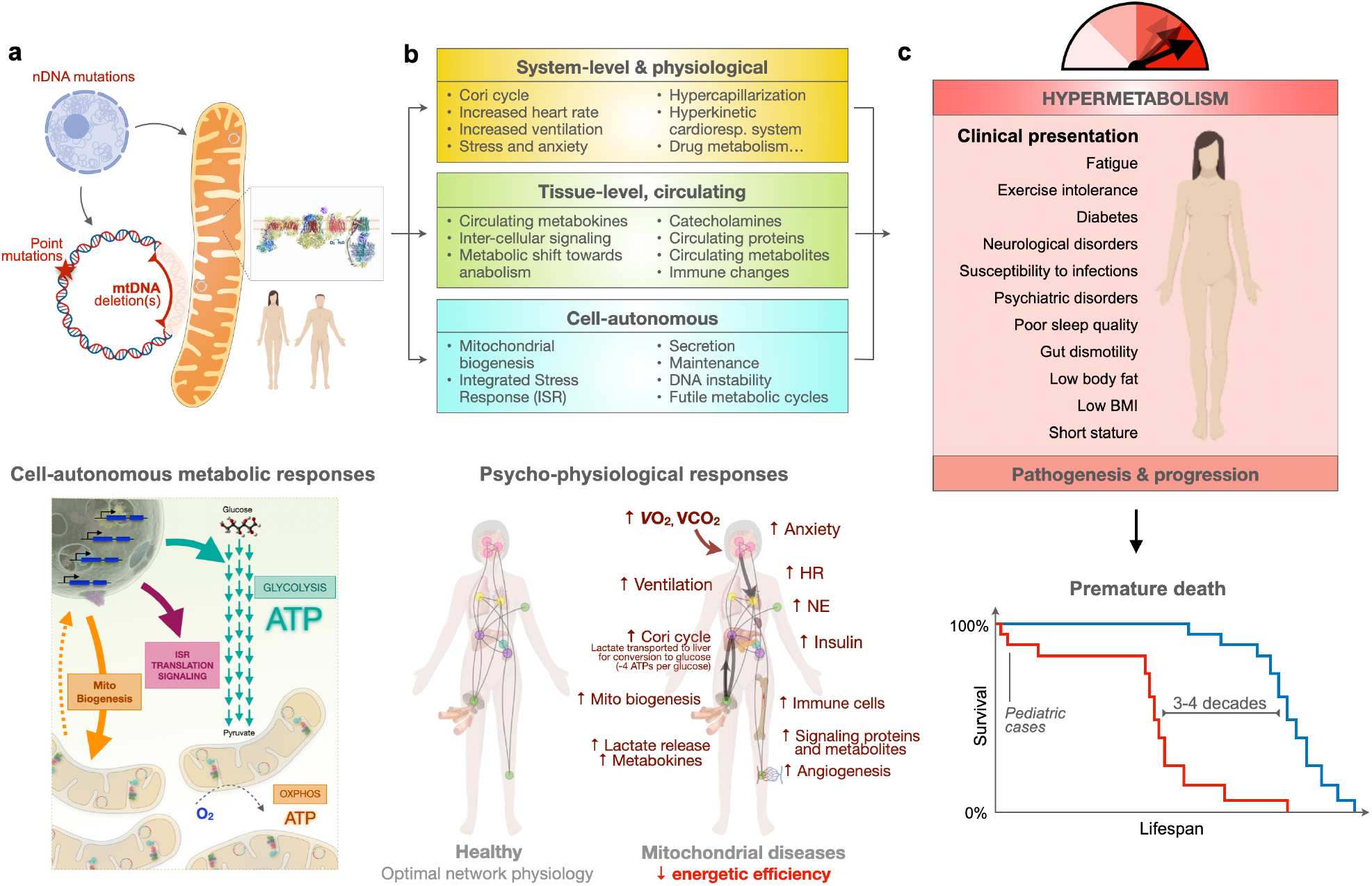
MulD-level recalibrations and documented compensatory adaptations and stress responses in patients with OxPhos defects. **a**) Genetic defects in the mitochondrial (mtDNA) or nuclear (nDNA) genomes cause OxPhos defects, which steer metabolic fluxes away from OxPhos and towards glycolysis in affected cells. **b**) OxPhos defects trigger physiological systems-level, tissue level, and cell-autonomous stress (i.e., compensatory) responses. The energetic cost of canonical mitochondrial disease features – e.g., mitochondrial hyperproliferation in ragged red fibers, the integrated stress response (ISR) and synthesis and release of metabokines (e.g., GDF15, FGF21), elevated catecholamines – has been unappreciated. These processes entail metabolic demands and culminate in hypermetabolism. **c**) As a result, the physiological effects of hypermetabolism and energy constraints on physiological and organ systems account for the clinical manifestations and pathogenesis of mitochondrial diseases, which ultimately shorten lifespan.

Further clinical studies using complimentary state-of-the-art EE assessments^26,34,35^ are needed to determine which subsets of individuals with genetic OxPhos defects are hypermetabolic, and when. Understanding the source of hypermetabolism, including cellular, tissue-level, and systemic recalibrations that contribute to EE also appear important to define. If hypermetabolism is a consistent clinical feature in this population, studies also will be needed to understand how hypermetabolism may contribute to symptoms and disease progression, and to develop strategies to reverse or mitigate its effects.

## Supplemental Material

### Methods

*Animal study meta-analysis*. Pubmed was searched using the following keywords: “mitochondrial disease”, “mitochondrial disorder”, “mitochondrial mutation”, “energy expenditure”, and “obesity”. Articles were also retrieved by manual curation. Data was extracted direction from tables or graphs, and meta-analyzed from 29 animal mitochondrial mutation model cohorts. Inclusion criteria included (1) study populations with control animals and experimental groups with defined genetic mitochondrial defects and (2) quantitative measurement of EE. Effect sizes of change in EE and physical activity were computed as percent change relative to control and significance was extracted from the primary publications. P-values computed by chi square.

*Human m*.*3243A>G mtDNA mutation cohort re-analysis*. Data of a controlled human cohort with the m.3243A>G mtDNA mutation were re-analyzed from Sharma et al., 2021^25^. The number of differentially represented plasma proteins and metabolites were kindly provided by the authors and p-values were computed using the Chi square test. Effect size for differences in clinical blood measures were calculated as percent change relative to control, and p-values were extracted from the original publication, computed by Wilcoxon ranked sum test.

